# Pharmacological Modulation of Autophagy Prevents Mutant SOD1^G93A^ Induced Neurotoxicity in Experimental Models of Amyotrophic Lateral Sclerosis (ALS)

**DOI:** 10.1101/2022.11.02.514252

**Authors:** Yue Bi, Xin Yan, Nazmiye Yacipi, Weisong Duan, Chunyan Li, Martin Thompson, Zhiying Shan, Lanrong Bi

## Abstract

Amyotrophic lateral sclerosis (ALS) is a fatal neurodegenerative disorder resulting from the progressive loss of both upper and lower motor neurons in the cerebral cortex, brainstem, and spinal cord. Currently, there are only two drugs, Riluzole (Rilutek) and Edaravone (Radicava), approved by FDA for ALS treatment. These two drugs are very expensive with only a few months of life extension. So far, there is no cure for ALS. Aberrant protein aggregation in motor neurons is an intracellular hallmark of ALS. The disturbance in protein homeostasis may contribute to the onset and progression of ALS. Autophagy plays an important role in degrading misfolded proteins, thereby preventing their aggregation. Pharmacological manipulation of autophagy has been proposed as a new therapeutic approach for treating ALS. IADB, a novel indole alkaloid derivative, has been reported to exert mitochondrial protection and cardioprotection through its autophagy-modulating potential. Our present study attempted to examine whether IADB has therapeutic potential in SOD1^G93A^-associated experimental models of ALS. We found that IADB could promote the clearance of SOD1^G93A^ aggregates and reduce the overproduction of mitochondrial reactive oxygen species (mtROS) in motor neuron-like NSC-34 cells transfected with SOD1^G93A^. We further examined the IADB in a SOD1-G93A mouse model of ALS. Administration of IADB started at the age of 55 days until the end stage of the disease. IADB treatment significantly increased LC3-II levels and decreased human SOD1 levels and p62 expression in the spinal cords of SOD1^G93A^ mice, suggesting that IADB treatment could induce autophagy activation and promote clearance of mutant SOD1 aggregates in this mouse model of ALS. Moreover, IADB treatment could alleviate the activation of microglia and astrocytes and reduce mitochondrial oxidative damage in the spinal cord of SOD1^G93A^ mice. Finally, we demonstrated that IADB treatment could improve motor performance and delay the onset and progression of the disease in a mouse model of ALS. The neuroprotective effects of IADB may mainly originate from its autophagy-promoting property.

## Introduction

Amyotrophic lateral sclerosis (ALS) is one of the leading causes of death by neurodegenerative diseases. The pathological feature of ALS is the progressive loss of both upper and lower motor neurons.^1^ ALS patients experience gradual paralysis of the voluntary muscles and ultimately death due to respiratory failure within 2-5 years of symptom onset. Currently, there are only two drugs, Riluzole (Rilutek) and Edaravone (Radicava), approved by FDA for ALS treatment. Riluzole can reduce excitotoxicity through the blockage of glutamatergic neurotransmission. ^2,3^ Edaravone can eliminate free radicals thereby reducing oxidative damage. ^2,4^ These two drugs are very expensive with only a few months of life extension. So far, there is no cure for ALS.

The major mechanisms of neurodegeneration in ALS proposed in the past decades include, but are not limited to, the following pathways: oxidative damage,^5^ abnormal protein aggregation,^6-8^ glutamate-induced excitotoxicity,^9,10^ neuroinflammation,^11,12^ mitochondrial dysfunction^13-16^, and RNA dysregulation ^17-19^. Aberrant protein aggregation in motor neurons is an intracellular hallmark of ALS.^20^ Intracellular accumulations of dysfunctional organelles and mutant proteins indicate the impairment in protein quality control pathways. Protein aggregates and damaged organelles can be trapped and degraded through the autophagic pathway. Autophagy is a highly conserved eukaryotic pathway for sustaining cellular homeostasis through the degradation of damaged intracellular materials.

The accumulation of protein aggregates is a typical sign of motor neuron degeneration in ALS.^21^ The disturbance in protein homeostasis may contribute to the onset and progression of ALS. Pharmacological manipulation of autophagy has been proposed as a new therapeutic approach for treating ALS. Mutations in the Cu, Zn-superoxide dismutase (SOD1) gene are known to contribute to the development and progression of ALS.^22^ Great research efforts have targeted the ALS-linked mutated SOD1 protein. In recent years, antisense therapy has been developed for the treatment of ALS. For example, Tofersen (BIIB067), a specific DNA fragment designed to target mutated SOD1 mRNA and promote its degradation, is in Phase III clinical trials.^23,24^ Niwa et al. have shown that the overexpression of Dorfin (a RING finger-type E3) ubiquitin ligase) could reduce SOD1 inclusions and protect neurons against neurotoxicity.^25^ Wada et al. have reported that induction of autophagy could decrease mutant SOD1 protein levels and accordingly, reduce their associated neurotoxicity.^26^ These data suggest that either autophagy enhancers or activators of the ubiquitin-proteasome pathway may have the therapeutically potential for the treatment of ALS.

Nitroxide derivatives can attenuate oxidative damage in various experimental models of oxidative stress-mediated diseases.^35,36^ The protective effects of nitroxides are attributed to their free radical scavenging capacities. Nitroxide also attenuates the formation of other reactive oxygen and nitrogen species. Unlike antioxidants that act in a sacrificial mode, nitroxides can provide a type of catalytic protection. Nitroxides undergo one-electron redox reactions to yield the corresponding hydroxyl-amines and oxoammonium cations via electron transfer reactions.^37^ The hydroxylamine and oxo-ammonium cations can comproportionate, yielding two nitroxide molecules. The non-radical species might also com-proportionate and yield the more stable radical form, thereby replenishing themselves. As a result, three forms of nitroxide derivatives (nitroxide radical, oxo-ammonium cation, hydroxylamine) can be present in the tissue. ^35-37^ Through the continuous exchange, the three forms can act as self-replenishing antioxidants, thereby bestowing catalytic protective activity. This key feature implicates the potential of this unique class of radical scavengers against oxidative damage. Tempol (4-hydroxy-Tempo), a cyclic nitroxide, is known to have excellent free radical scavenging activity and cellular permeability. Neuroprotective effects of Tempol have been reported in various diseases models, such as ischemic stroke,^38-40^ ischemia-reperfusion injury,^40-45^ and neurodegenerative diseases including ALS.^46-49^

Indole alkaloids are naturally occurring plant substances that have a wide spectrum of neuropharmacological,^27^ psychopharmacological, ^27,28,^ and antioxidants effects.^29^ It has been reported that a tryptophan microbiota metabolite functions as a regulator of autophagy and may have therapeutic implications in inflammatory and autophagy-mediated disorders.^30^ Indole-3-carbinol (I3C) has been found to promote autophagy in hyperlipidemia zebrafish larvae.^31^ Recently, it has been reported that I3C combined with hydroxychloroquine (HCQ) exhibited better antitumor activities than each drug alone via targeting autophagy and apoptosis.^32^ 3,3’-diindolylmethane (DIM) is a compound found in the broccoli family and it has been reported to induce autophagy and apoptosis in some types of human cancer.^33^ In addition, DIM has been reported to attenuate inflammation and apoptosis induced by cardiomyocyte hypoxia via activation of autophagy. ^34^

The indole derivatives have been reported to possess anti-inflammatory, free radical scavenging activity, mitochondrial protection, and autophagy-modulating activity.^50-55^ For example, IADB, a novel indole alkaloid-nitroxide conjugate, has been reported to exert mitochondrial protection and cardio-protection through its autophagy-modulating potential.^52^ In our present study, we attempted to further examine the role of IADB in the SOD1^G93A^-associated experimental models of ALS.

## Materials and Methods

### Cell culture and transfection

The mouse motor neuron-like NSC-34 is a hybrid cell line produced by the fusion of motor neurons from the spinal cord embryos with N18TG2 neuroblastoma cells. NSC34 cells were grown in Dulbecco’s modified Eagle’s medium (DMEM) supplemented with 10% heat-inactivated fetal bovine serum (FBS), 100 IU/mL penicillin, and 100 µg/mL streptomycin at 37°C in a 5% CO_2_ incubator. NSC-34 cells were transiently transfected with a mutated human SOD1 (G93A) cDNA fused to an enhanced green fluorescent protein (eGFP) or empty vector (EV), respectively, using Lipofectamine 2000 following the manufacturer’s instructions. To examine the effect of IADB on mutant SOD1, IADB (30 µM) was added to the growth medium of transiently transfected NSC-SOD^G93A^ or NSC-EV cells and then incubated for 24h. The effects of IADB on the expression of mutant SOD1 and the autophagy markers (LC3-II and p62) in NSC34 cells were examined by Western blot.

### Animal model

Transgenic mice B6SJL-TGN (SODG93A) were originally obtained from the Jackson Laboratory. This strain of mice expresses the mutant human SOD1 gene which harbors a single amino acid substitution of glycine to alanine at codon 93. The mice gradually exhibit a phenotype similar to ALS in humans; becoming paralyzed in one or more limbs with paralysis due to loss of motor neurons from the spinal cord. A colony of SODG93A was maintained by breeding the SODG93A mice (B6SJL-TGN (SOD^G93A^) 1Gur/J, the Jackson Laboratory) with non-transgenic (B6SJLF1 mice, the Jackson Laboratory). Mouse DNA was extracted to determine the genotypes of transgenic mice by polymerase chain reaction (PCR) as described previously. Non-transgenic (NTG) littermates not expressing the SOD1^G93A^ were used as wild-type controls. Animals were housed under standard housing conditions with a 12-h light/dark cycle, and food and water *ad labium*. The animal protocols proceeded by the Guidelines for Preclinical Animal Research in ALS/MND.^56^

IADB (40mg/kg) was administrated intraperitoneally daily starting at 55 days of age in both SOD1^G93A^ mice and NTG littermates. The animals in the NTG control group received an equal volume of normal saline instead of IADB treatment. After 60 days of treatment, part of the mice was sacrificed for pathological examination and the other part of the animals were continuedly treated until the end stage. Male SOD1^G93A^ mice were randomly assigned into the following groups: NTG; NTG + IADB; SOD1^G93A^; SOD1^G93A^ + IADB) with 14 mice in each group. To monitor disease progression, animals were examined daily for checking motor deficits. The body weight was measured every three days, starting at the age of 70 days until the end stage. The neurological score (0-4) was defined following the ALS Therapy Development Institute. ^69^

#### Rotarod test

the motor function was evaluated following the published protocol.^70^ In brief, mice were trained for 3 days, and then basal motor performance was recorded. The disease onset was defined under the condition that one of the following 3 criteria is fulfilled: (1) abnormal gait is detected; (2) unable to stay on the rod for 5 min; (3) two consecutive instances of weight loss are observed.

#### Evaluation of life span

the animals that are unable to right themselves within the 30s after being placed on either side of the back were defined as the end-stage.

### Immunofluorescence staining

Mice were deeply anesthetized with chloral hydrate and then were transcardially perfused with ice-cold phosphate-buffered saline (PBS, pH 7.4). The spinal cords were collected and fixed in 4% paraformaldehyde for 24h at 4°C. Then the spinal cords were transferred and kept in 30% sucrose in PBS at 4°C until they sank to the bottom. Serial frozen sections of the spinal cord were cut and mounted on gelatin-coated slides. The sections from the animals of different treatment groups were washed using PBS and permeabilized with 0.3% Triton X-100 and followed by incubation overnight at 4°C with the following primary antibodies:: rabbit anti-NeuN (1:100, Sigma-Aldrich), mouse anti-GFAP (1:500, Millipore), rabbit anti-IBA-1 (Wako chemicals), LC3B (1:200; Cell Signal), SOD1 (1:200; Abcam), SQSTM1 (1:100; BD Transduction Laboratories). Then the slides were incubated with the corresponding secondary antibody for 2h.

### Statistical analysis

One-way ANOVA was used for the analysis of the data of immunofluorescence and qRT-PCR. The results of the rotarod test and body weight were analyzed using two-way ANOVA for repeated measures. The level of statistical significance was 95% (p<0.05).

## Results

### IADB treatment attenuated mutant SOD1^G93A^-induced oxidative stress in NSC-34 motor neuron-like cell lines

The mouse motor neuron-like NSC-34 cell line is a hybrid of the embryonic mouse spinal cord motor neuron and the neuroblastoma cell line. NSC-34 cells have the key features of motor neurons, such as the extension of neurites, generation of action potentials, expression of neurofilament proteins, and so on. ^57^ As excess oxidative stress has been observed in the motor neurons of ALS patients,^58^ we attempted to examine the mitochondrial oxidative status in NSC-34 cells transfected with empty vector (EV) or mutant SOD1^G93A^. Toward this, the transfected NSC-34 cells were labeled with MitoProbe,^64^ a specific mitochondrial-targeted fluorescent probe for the detection of reactive oxygen species (ROS) generation in the mitochondria (mt). The mtROS generation was further confirmed in a large population of NSC-34 cells by flow cytometry. Compared to the empty vector-transfected cells, SOD1^G93A^ transfected NSC-34 cells showed an approximate ∼2.8-fold increase in fluorescence emission from MitoProbe, suggesting enhanced generation of mtROS occurring in NSC cells expressing mutant SOD1^G93A^ (**Figure 1A**). Compared to the cells surrounding the EV-transfected NSC cells, a significant increase in terms of mtROS generation was also observed in neighboring cells of NSC-SOD1^G93A^, implying the vulnerability of NSC cells to the neurotoxicity of mutant SOD1^G93A^ protein. Interestingly, the mitochondrial oxidative damage observed in the SOD1^G93A^ transfected cells, and their neighboring cells could be significantly inhibited when cells were treated with IADB (**Figure 1A**).

**Figure 1.**
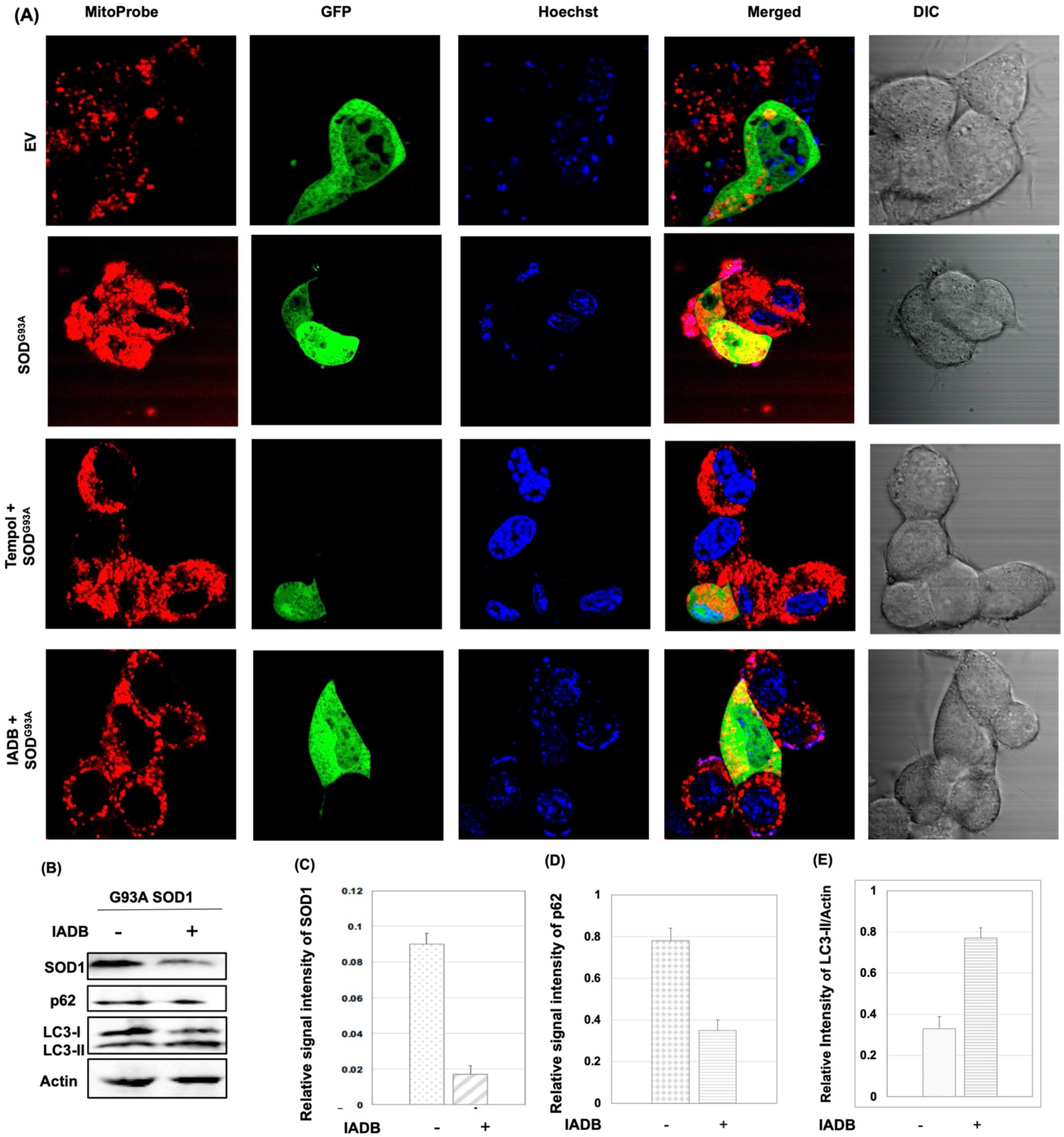
IADB treatment reduces the aggregates of mutant SOD1 in motor neuron-like NSC 34 cells thereby protecting neuronal cells against mitochondrial oxidative damage. (**A**) Representative confocal microscopic image of NSC-34 cells transfected with empty vector (EV) or GFP-SOD1^G93A^ (green fluorescence) and then counter-stained with MitoProbe (red fluorescence), and Hoechst 33342 (blue fluorescence): Fluorescence images are displayed for cells: NSC-EV (1^st^ row); NSC-SOD1^G93A^ (2^nd^ row); NSC-SOD1^G93A^ + Tempol (30 µM) (3^rd^ row); NSC-SOD1^G93A^ + IADB (30 µM) (4^th^ row). (**B**)-(**E**): IADB treatment reduces the expression of mutant SOD1 and induces autophagy activation in NSC-34 cells transfected with SOD^G93A^. Western blot analysis (**C**) shows the expression of SOD1, p62, and LC3-I/II in NSC-34 cells transfected with mutant SOD^G93A^ before/after IADB treatment. The relative levels of SOD1 (**C**), p62 (**D**), and LC3-II (**E**) were quantified by densitometry. β-actin was used as a loading control. Error bars indicate ± SEM from at least four independent experiments. Data were presented as the mean ± SD (n=3).

### IADB treatment reduced levels of mutant SOD1 via activation of autophagy in motor neuron-like NSC-34 cell lines

To examine the effect of IADB on mutant SOD1, IADB (30 µM) was added to the growth medium of NSC-SOD1^G93A^ cells. After 24 h incubation with the medium containing IADB, the levels of mutant SOD1 in NSC-34 cells were significantly reduced (**Figure 1B-E**). It is known that autophagic activity is important for preventing the accumulation of abnormal proteins, we then compared the autophagic response of NSC-SOD1^G93A^ cells before and after IADB treatment. Proteins light chain 3 (LC3) and p62 are correlated with the formation of autophagosomes during which cytoplasmic contents are engulfed for subsequent degradation. We found that the level of LC3-II increased significantly accompanied by a noticeable decrease in p62 in IADB-treated NSC-SOD1^G93A^ cells (**Figure 1B-E**). These data suggest that IADB could induce autophagy activation in NSC-SOD1^G93A^ cells.

### IADB treatment stimulates activation of autophagy in the SOD1^G93A^ mouse model of ALS

Mutation in the sequestosome 1 (SQSTM1) gene which encodes the p62 protein has been identified in familial and sporadic cases of ALS. ^59^ P62 is a multifunctional protein involved in protein degradation both through proteasomal regulation and autophagy.^60^ Many ALS patients have mutated p62 protein, and dysfunction of P62 contributes to impairment in the protein degradation system which removes misfolded and damaged proteins.^60-62^ Therefore, the expression of the p62 protein is used as a marker of autophagy, which is degraded with the activation of autophagy.^60-63^ IADB treatment significantly increased LC3-II levels but decreased human SOD1 (hSOD1) levels and p62 expression in the spinal cords taken from IADB + SOD1^G93A^ and IADB + NTG mice (**Figure 2**), suggesting that IADB treatment could induce autophagy activation and promote clearance of mutant SOD1 aggregates in the SOD1^G93A^ mouse model of ALS.

**Figure 2.**
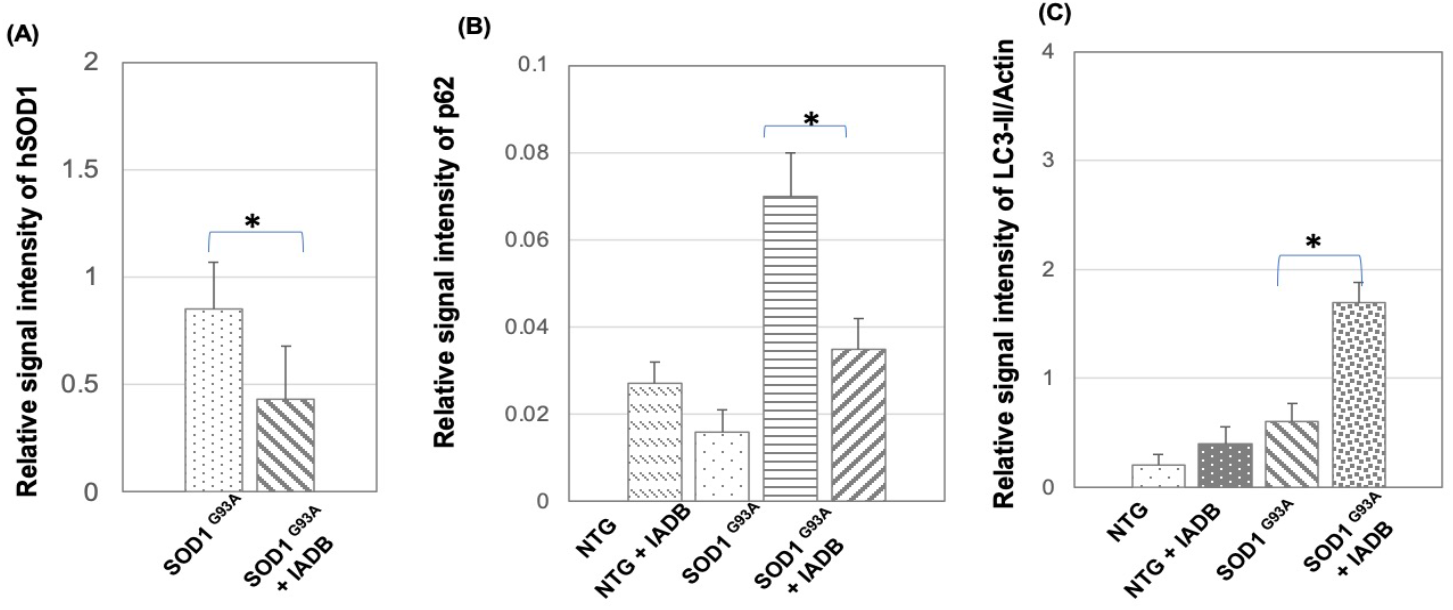
IADB treatment reduces the expression of mutant SOD1 and stimulates autophagy activation in SOD1^G93A^ mice. Western blot analysis of the protein levels of mutant human SOD1 (hSOD1) (**A**), the protein levels of p62 (**B**), and LC3-I/II (**C**) in the spinal cords of SOD1^G93A^ mice were quantified and compared with/without IADB treatment.

### Induction of autophagy attenuates microglial and astrocytic activation and mitochondrial oxidative damage in the spinal cord of SOD1^G93A^ mice

To further examine the effect of IADB on the mitochondrial oxidative status in *in vivo*, MitoProbe (30 µM) was intraperitoneally injected in NTG- and SOD^G93A^-mice, respectively.^64^ mtROS production was estimated based on the measurement of the mean fluorescence intensity (MFI) of MitoProbe staining. The SOD-G93A mice (115 days old) showed a greater number of MitoProbe-positive granules with stronger red density compared to the NTG-mice (115 days old). We found that the IADB treatment resulted in a significant decrease in terms of the number and the fluorescent intensity of MitoProbe-positive granules in microglial cells when compared to that in the groups of SOD1^G93A^ mice. To determine the effect of IADB on microglial cells, we performed Iba1 immunostaining to examine microglial cells in the spinal cord of SOD1^G93A^ mice. We found there was a significantly higher number of Iba1 positive microglial cells in the spinal cord of SOD1^G93A^ mice compared with that in NTG mice (**Figure 3A-C**, p<0.01), suggesting strong microgliosis. Interestingly, we observed that IADB-treated SOD1^G93A^ mice showed a much lower number of activated microglial cells than that of the control group of SOD1^G93A^ mice (**Figure 3A-C**, p<0.01). However, there was no statistical difference in terms of Iba1-positive microglial cells between the IADB-treated SOD^G93A^ mice and NTG mice.

**Figure 3.**
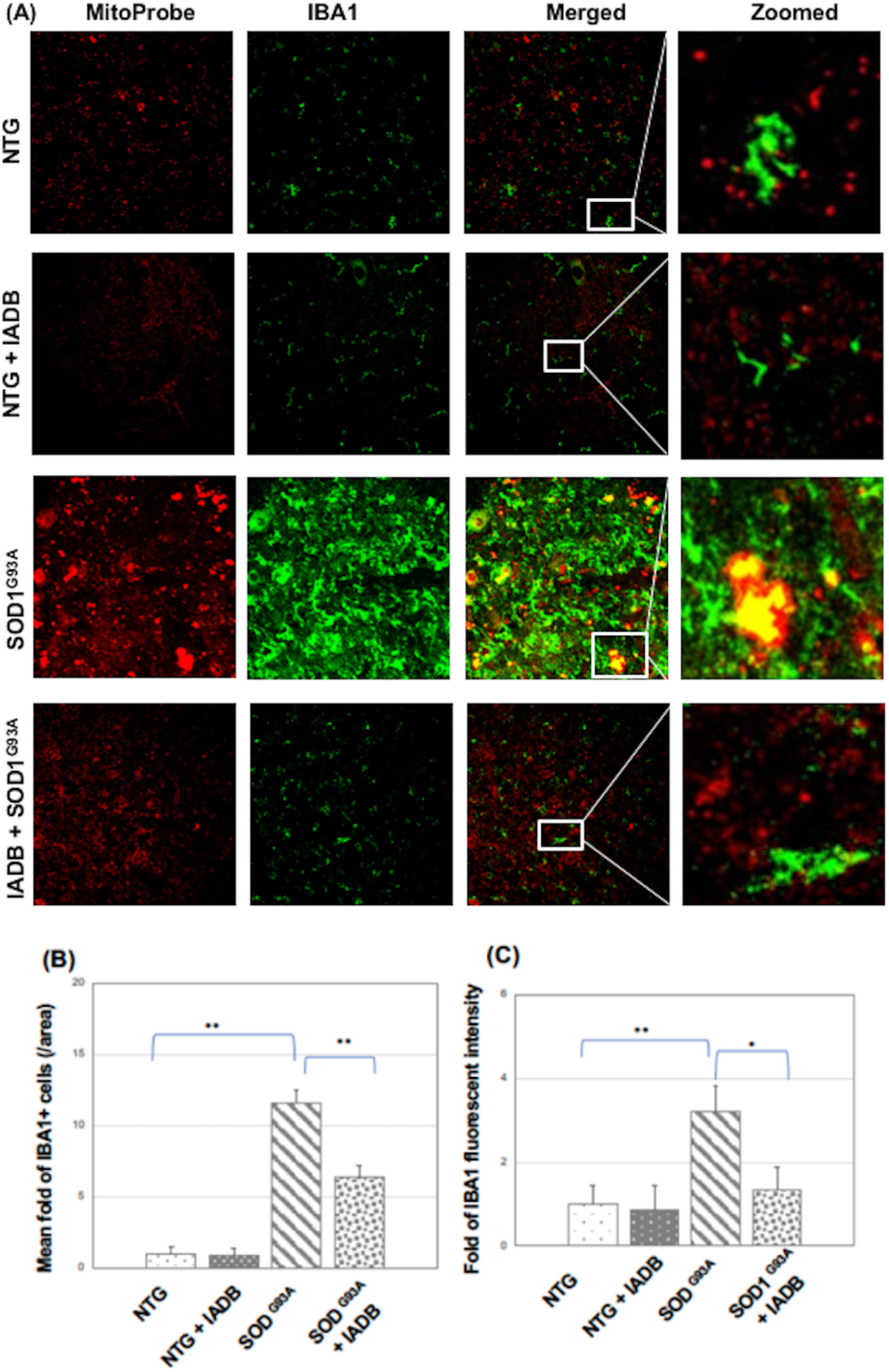
IADB treatment attenuated microgliosis in the lumbar anterior horn of SOD1^G93A^ mice. Iba1 immunostaining labeling (green fluorescence) activated microglia in the lumbar anterior horn of four different groups of mice at the age of 115 days, and the samples were counterstained with MitoProbe (red fluorescence) for detection of mitochondrial ROS generation (**A**). Relative changes in microglia numbers are expressed as numbers per unit of area (**B**) and fluorescence intensity of Iba1 immunostaining (**C**).

We further examined the effect of IADB on the action and ROS production of astrocytes in the spinal cord of SOD1^G93A^ mice and its normal controls. GFAP (a marker for astrocytes) immunostaining was performed using the spinal cord sections, We found that there was a significantly higher number of GFAP-positive astrocytes in the spinal cord of SOD1^G93A^ mice compared with that in NTG mice (115 d) (**Figure 4A-C**, p<0.01), suggesting strong astrogliosis. It was observed that IADB-treated SOD1^G93A^ mice (115 d) showed a much lower number of activated astrocytes than that in the control group of SOD^G93A^ mice (115 d) (**Figure 4A-C**, p<0.01). No statistical difference was observed in GFAP-positive reactive astrocytes between the IADB-treated SOD1^G93A^ mice and NTG littermate. Similarly, we noticed that the IADB treatment resulted in a significant decrease in the levels of mtROS in astrocytes when compared to that in the control group of SOD1^G93A^ mice.

**Figure 4.**
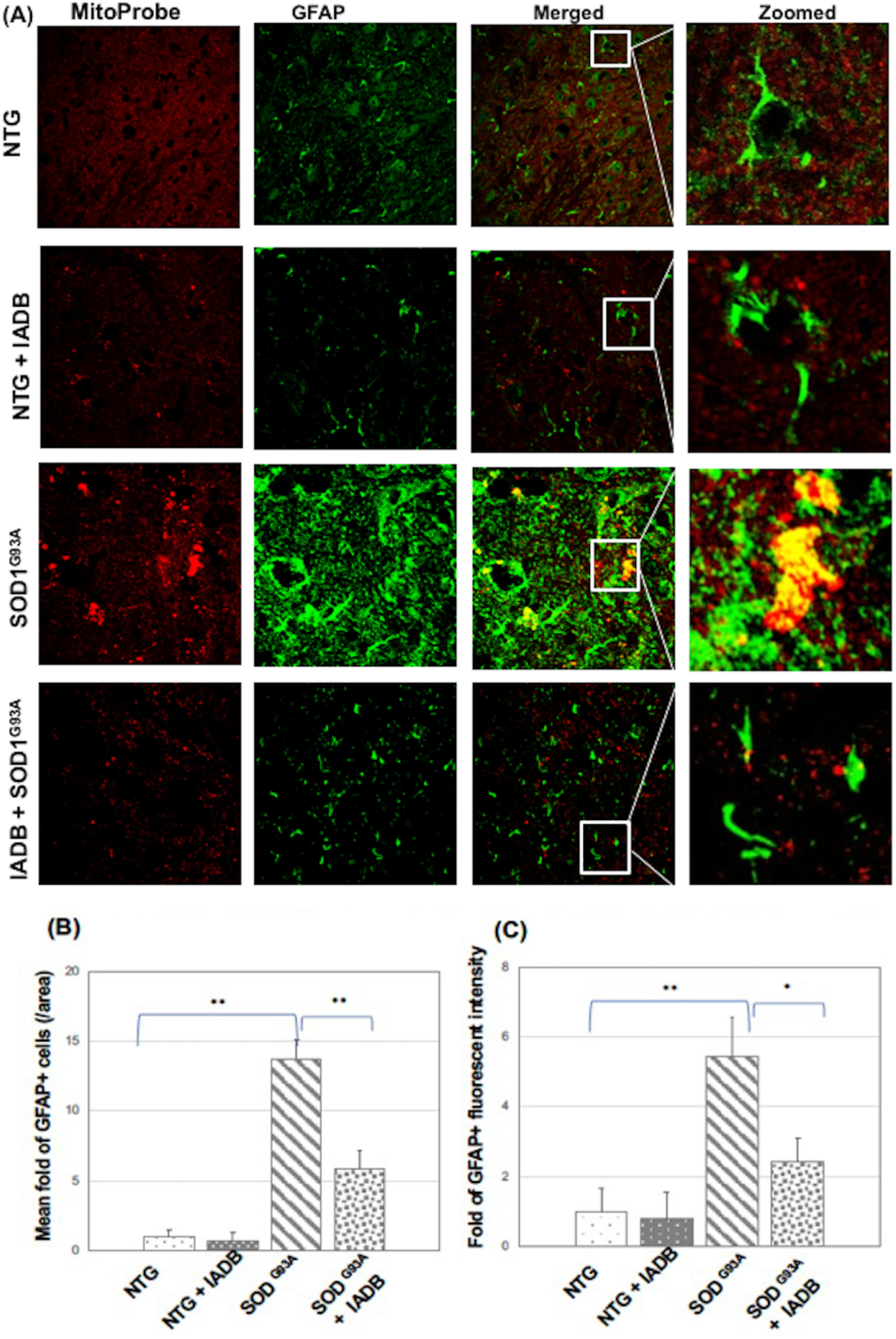
IADB treatment attenuated astrogliosis in the lumbar anterior horn of SOD1^G93A^ mice. GFAP immunostaining labeling (green fluorescence) activated astrocytes in the lumbar anterior horn of four different groups of mice at the age of 115 days, and the samples were counterstained with MitoProbe (red fluorescence) for detection of mitochondrial ROS generation (**A**). Relative changes in activated astrocytes are expressed as (**B**) numbers per unit of area and (**C**) fluorescence intensity of GFAP immunostaining.

### Induction of autophagy could delay disease onset and progression and improve survival in SOD1^G93A^ mice

To examine whether IADB treatment can affect disease onset and progression, we performed the rotarod test in the SOD1^G93A^ mice for evaluating motor function. It was noticed that animals from IADB + SOD1^G93A^ group exhibit significantly delayed disease onset compared to those from the SOD1^G93A^ group (114.0 ± 1.5 *vs* 92.0 ± 1.0 d, p<0.01). Animals from IADB + SOD1^G93A^ group showed a noticeable life extension compared with those from SOD1^G93A^ treated with vehicle (148.0 ± 2.1 vs 122.1 ± 1.5 d, p<0.01). Animals from Tempol + SOD1^G93A^ group showed no significant change in survival compared to SOD1^G93A^ treated with a vehicle.

## Discussion

ALS is characterized by progressive loss of upper and lower motor neurons in the spinal cord, brainstem, and motor cortex.^1^ The mechanisms underlying ALS development remain elusive. Growing evidence suggests that oxidative damage may be responsible for the onset and/or progression of ALS.^5,13,16^ Elevated levels of ROS have been proposed to cause mitochondrial DNA mutations, which are frequently observed in the motor cortex and spinal cord of ALS patients.^65^ Previous studies pointed out that the increased production of ROS is an early and likely causative event that may eventually lead to neuronal death. ^65^ Mitochondria are the major site of ROS production, and they are also targets of ROS. Mitochondrial dysfunction caused by oxidative damage is considered a major cause of aging. Morphologically abnormal mitochondria have been often observed in the motor neurons of ALS patients. ^14-16^ It has long been debated whether oxidative stress is a primary cause of pathogenesis in ALS or is just a consequence of the disease. Low levels of ROS are important messengers in many cellular processes including cell growth, differentiation, and inflammation.^13^ Under normal conditions, the production, and removal of ROS can be balanced by the cellular antioxidant defense system. Excessive ROS production associated with an inefficient antioxidant defense represents an important pathological feature in ALS. ^5,13,16^ The presence of SOD1^G93A^ in the mouse model of ALS often leads to a shift in the mitochondrial redox balance toward a higher level of mtROS.^5^ Although the antioxidant strategy is a very promising approach to slow the progression of ALS, SOD^G93A^ mice treated with an antioxidant, Tempol, did not extend the lifespan of SOD^G93A^ mice compared to SOD^G93A^ mice treated with vehicle, suggesting that preventing peroxide-mediated mitochondrial damage alone is not sufficient to delay the diseases.

The aggregation of misfolded proteins is frequently observed in the motor neurons of both sporadic ALS and familial ALS patients. ^6-8^ Therefore, it is reasonable to propose that drugs that are capable of either preventing the aggregation of misfolded proteins or promoting its degradation should be able to slow the onset and progression of ALS. The indole derivatives have been reported to possess anti-inflammatory, free radical scavenging, mitochondrial protection, and autophagy-modulating activity. ^50-55^ In our present study, we attempted to examine the effect of a novel indole derivative, IADB, upon the SOD1^G93A^-related neurotoxicity in the experimental models of ALS.

Autophagy is a catabolic pathway in which dysfunctional cellular components can be degraded via the lysosome and are recycled. Autophagy is a multi-step process including the formation of autophagosomes and autolysosomes (the fusion of the autophagosome with the lysosome) followed by the degradation of the contents of the autolysosome.^8^ The microtubule-associated protein 1A/1B-LC3 and the autophagy receptor SQSTM1 (p62) are commonly used autophagy markers.^59-63^ During autophagy, cytosolic LC3-1 is conjugated to phosphatidylethanolamine to form LC3-II; LC3-II is then incorporated into the autophagosomal membrane. The conversion from LC3-I into LC3-II is accepted as a simple method for monitoring autophagy. ^8^

The p62 protein is a receptor for the delivery of cargo for degradation via autophagy, which can bind ubiquitin and LC3, thereby assisting the clearance of ubiquitinated proteins. p62 protein is a commonly used marker for monitoring autophagy, which is degraded with the activation of autophagy. ^59-63^ We have demonstrated that IADB treatment could upregulate LC3 and downregulate p62 in a cellular model of ALS (**Figure 1**). Therefore, it is reasonable to assume that IADB has a neuroprotective effect against SOD1^G93A^ -induced neurotoxicity and alleviates mitochondrial oxidative damage through the induction of autophagy. Next, we further examined the effect of IADB in an animal model of ALS. Daily IADB treatment starts at the age of 55 days until the end stage of the disease. After IADB treatment, we noticed that the protein level of p62 decreased together with increasing protein levels of LC3 in the spinal cord of SOD1^G93A^ mice (**Figure 2B&C**). Induction of autophagy induced by IADB further led to reduced mutant SOD1 levels in the spinal cord of SOD1^G93A^ mice (**Figure 2A**).

Microglia are the first responders of immune defense in the central nervous system. They constantly examine the surrounding microenvironment and respond to cellular distress. ^66^ Under normal physiological conditions, microglia remain in an inactive state. Misfolded SOD1 aggregates formed pore-like structures in the lipid membrane resulting in the influx of calcium and microglia activation. ^66^ Chronic productions of inflammatory mediators, particularly the generation of ROS and •NO by reactive microglia not only regulate microglial activity but also impact motor neurons. During the initial phase of neuroinflammation, O_2_ ^• -^ is formed through the microglial NADPH oxidase system. As inflammation progresses to the chronic state, iNOS produces •NO. O_2_ ^• -^ reacts with •NO radical produces ONOO^−^. ONOO^−^ reacts with CO_2_ to generate a series of highly reactive radicals, such as •NO_2_, •OH, and •CO_3_. Each of them can initiate lipid peroxidation. The SOD1 mutation reduces the cell’s ability to remove harmful free radicals, thereby further aggravating lipid peroxidation. Once lipid peroxidation is initiated, an excess of highly reactive lipid peroxidation end products is produced (i.e., acrolein (ACR), 4-hydroxy-2-nominal (HNE), and malondialdehyde (MDA)).^67,68^ Accumulation of peroxidated lipids led to neuroinflammation, demyelination, and neurodegeneration. For example, HNE levels were significantly elevated in the sera and spinal fluid of sporadic ALS patients compared with control populations and positively correlated with the extent of the disease. ^68^ Oxidative stress can activate glia cell-mediated inflammation which also causes secondary neural damage. In our present study, we found that IADB treatment could reduce elevated mitochondrial ROS and reactive glial cells in the spinal cord of SOD1^G93A^ mice (**Figure 3** and **Figure 4**). The autophagy-promoting property of IADB may contribute to the reduction of mitochondrial oxidative damage and glial activation (including microglia and astrocytes) in SOD1^G93A^ mice.

### Conclusion

In our present study, we have demonstrated that IADB could promote the clearance of SOD1^G93A^ aggregates and reduce the overproduction of mitochondrial ROS in a cellular model of ALS. The in vivo data obtained from SOD1^G93A^ mice are consistent with the results obtained from cellular assays. IADB treatment upregulated LC3 and downregulated p62 in the SOD1G93A-associated models of ALS. In addition, we have shown that IADB treatment could alleviate the activation of microglia and astrocytes in SOD1^G93A^ mice. Furthermore, we have demonstrated that IADB treatment improved motor performance and prolonged the survival of SOD1^G93A^ mice. Taken together, we have demonstrated that IADB could induce autophagy in SOD1^G93A^ associated cellular model and animal model of ALS. The enhancement of autophagy induced by IADB treatment leads to the reduction of the aggregation of mutant SOD1, and this is an important step to achieving neuroprotection.

